# Trophic transfer of nanoplastics reduces larval survival of marine fish more than waterborne exposure

**DOI:** 10.1101/2025.09.10.675286

**Authors:** Siti Syazwani Azmi, Ozan Oktay, Hee-Jin Kim, Hisayuki Nakatani, Mitsuharu Yagi

## Abstract

Microplastics (MPs) and nanoplastics (NPs) are widespread contaminants in marine environments and pose significant risks to aquatic organisms. However, physiological effects and survival consequences of different MP and NP exposure pathways during early developmental stages of marine fish remain poorly understood. We investigated effects of direct and indirect consumption of MPs and NPs by larvae of red sea bream (*Pagrus major*) using a controlled laboratory system. Larvae were exposed to fluorescently labeled polystyrene particles (3 µm and 0.2 µm) either directly from the water or indirectly via contaminated zooplankton prey. MPs and NPs were detected in digestive tracts of all exposed individuals, regardless of the exposure route. However, survival was significantly reduced in larvae that consumed NPs via rotifer predation, suggesting that ingestion of contaminated prey organisms may represent a greater hazard than direct uptake from water. In both groups, antioxidant enzymes, superoxide dismutase (SOD) and catalase (CAT) were elevated, suggesting induction of oxidative stress. Relative gene expression revealed greater upregulation of several stress and immune-related genes in larvae exposed to MPs via rotifer predation than via direct consumption. Our findings provide clear evidence that both MPs and NPs can alter physiological and molecular responses in marine fish larvae, especially via rotifer predation. This study suggests the importance of considering trophic interactions in ecotoxicological assessments of plastic pollution to address plastic bioavailability and toxicity during early life stages.

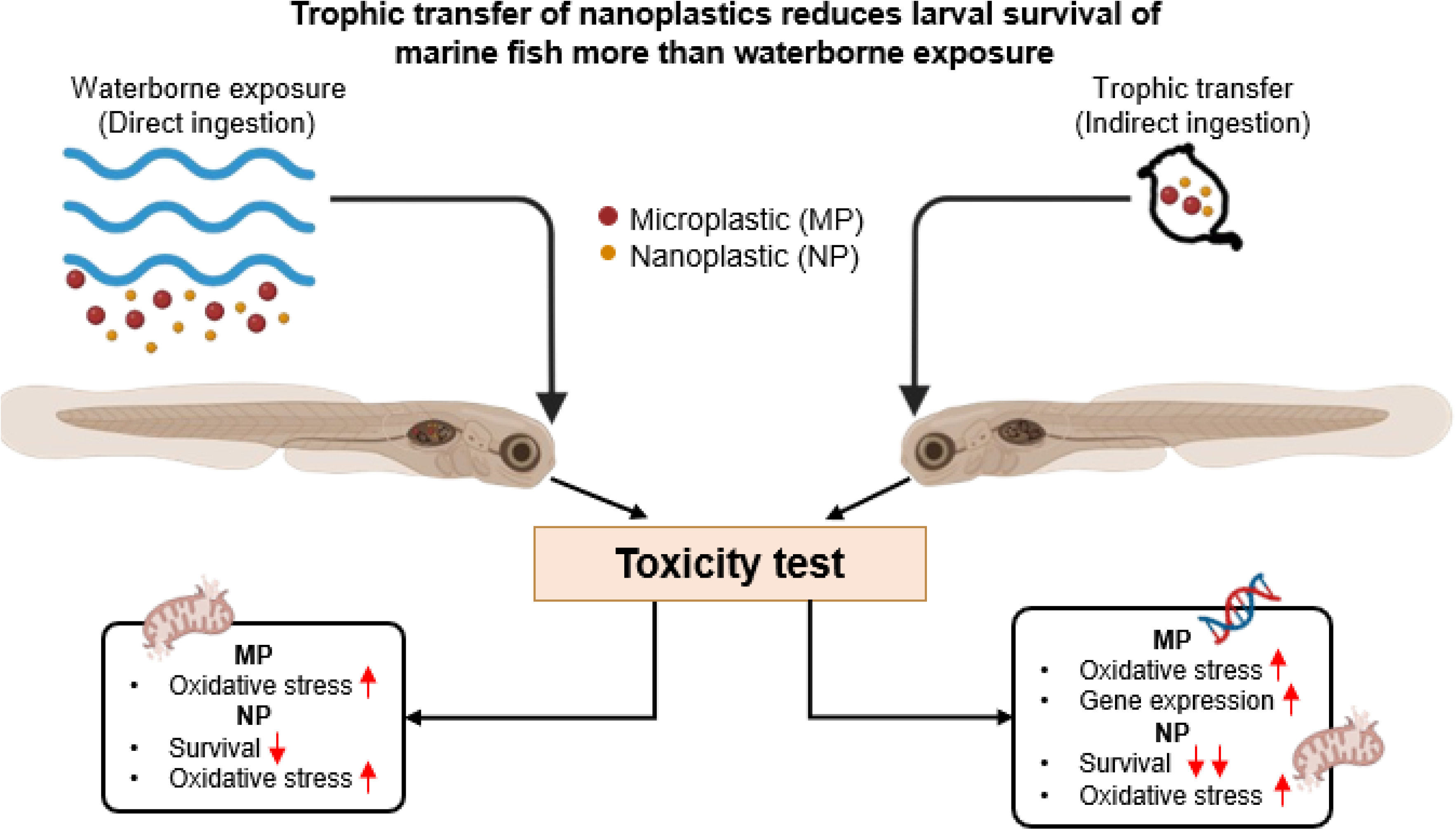

**Highlights:**

- Larval survival was significantly reduced by nanoplastics via trophic transfer
- Growth of larvae was not affected by micro-and nanoplastics
- Micro- and nanoplastics cause oxidative stress by both exposure routes
- Up-regulated gene expression revealed that larvae undergo various stresses

## 1. Introduction

The global increase of plastic pollution is driven by inadequate plastic waste management systems exacerbated by increasing plastic production and consumption. Global plastic production reached approximately 367 million tons in 2020 and continues to rise annually (Sajjad et al., 2022). This is the key driver increasing the plastic load in marine ecosystems. Although most plastic waste is recyclable, it is estimated that 5 – 12 million tons of plastic waste are released into aquatic environments every year (Bayhan & Uncumusaoglu, 2024). Once this plastic waste enters the environment, larger plastic pieces are degraded into smaller debris. These smaller plastic fragments, known as microplastics (MPs) measure < 5 mm. Moreover, further degradation results in formation of nanoplastics (NPs) ranging from 1 to 1000 nm (Kwak & An, 2021). Due to their small size and low biodegradability, they remain in the environment longer, contributing to plastic pollution and harming both plants and animals (Ding et al., 2018; Sendra et al., 2021; Feng et al., 2022).

MPs and NPs have emerged as critical environmental concerns, with a significant number of studies documenting plastic presence in various taxa (Santos et al., 2021; Abdel-Latif et al., 2022). Both MPs and NPs can be ingested by plankton (Carbery et al., 2018), fish (Lu et al., 2016; Ding et al., 2018; Feng et al., 2022; Yagi et al., 2022), bivalves (Abidli et al., 2023), birds (Lu et al., 2023; Navarro et al., 2023), and even mammals (Zantis et al., 2021). Ingestion and accumulation of both MPs and NPs have diverse adverse effects such as impaired growth, altered feeding and swimming behaviour, and blockage of the digestive system (Yang et al., 2020; Pelegrini et al., 2023). In addition, MP and NP ingestion can also reduce reproductive success and survival, alter physiological processes, cause organ damage, and trigger inflammatory responses and apoptosis (Lei et al., 2018; Le Bihanic et al., 2020; Chen et al., 2022). Fish consume MPs and NPs either by directly ingesting plastic pieces that they misidentify as food, or by consuming contaminated prey (Watts et al., 2014). The latter case, known as trophic transfer, may result in accumulation of both MPs and NPs in predators at higher trophic levels, potentially affecting the food chain (Carbery et al., 2018; Zhang et al., 2019). Accumulation of MPs and NPs in predators and effects of these plastics at higher trophic levels is still limited (Carbery et al., 2018; Lu et al., 2018).

In particular, there is little information available relative to biological impacts of microplastic ingestion on fish larvae, as most studies have focused on adult, freshwater fishes. Microplastic ingestion in freshwater larval fishes reduces hatching and survival, and impacts swimming behaviour, gene expression, and energy metabolism (Lu et al., 2016; LeMoine et al., 2018; Pannetier et al., 2020). Le Bihanic et al. (2020) reported that exposure to benzo(a)pyrene, perfluorooctanesulfonic acid and benzophenone-3 microplastics decrease embryonic survival and growth, reduce hatching, and increase developmental anomalies and abnormal behaviour of marine medaka after as little as 12 days. However, Campos et al. (2021) revealed limited impacts on survival and growth, but found several biochemical responses after 7 h of polyethylene MP exposure.

MPs and NPs, alone or in combination with other pollutants, also induce oxidative stress and cause abnormalities in lipid metabolism (Hamed et al., 2020; Yang et al., 2020). Production of excessive intracellular reactive oxygen species (ROS) is a well-recognized toxicity mechanism induced by chemical pollutants. ROS are mainly produced by innate immune cells in response to stressful xenobiotics (Yang et al., 2020). Neutrophils release ROS as a first defense against microbes, and antioxidant enzymes degrade low levels of ROS. However, excessive ROS production resulting from prolonged or intense pro-oxidant stimuli can overwhelm these defenses, disrupting redox signaling, and leading to oxidative damage (Hamed et al., 2020). That said, not all MPs and NPs activate antioxidant defenses, raising questions about their mechanisms of action. For example, Yang et al. (2020) observed increased activity of both glutathione peroxidase (GPx) and superoxide dismutase (SOD), and decreased activity of catalase (CAT) in goldfish (*Carassius auratus*) larvae exposed to polystyrene microsphere MPs or NPs for 7 days. Moreover, increased gene expression of SOD and CAT was also observed in zebrafish (*Danio rerio*) embryos exposed to polystyrene NPs for 24 h (Feng et al., 2022).

We assessed effects of MP and NP exposure on early-stage red sea bream (*Pagrus major*) larvae for 12 days through direct consumption and via predation on contaminated rotifers. We targeted early life stages of these fish because they serve as vital nutritional links between primary producers and higher predators. Few microplastic studies have targeted marine species, disruption of which could have substantial ecological consequences. In the current study, we examined impacts of MPs and NPs on red sea bream survival, growth, modulation of gene expression of SOD, CAT, glutathione S-transferase alpha 1 (GSTA1), heat shock protein (HSP70), and immune response genes [interleukin-8 (IL-8), interleukin-1 beta (IL-1β), and toll-like receptor (tlr)]. This species was chosen as an experimental model as it is a commercially important aquaculture species of high economic value (Hossain et al., 2016); however, there is no record of MP or NP exposure in red sea bream.

## 2. Materials and methods

### 2.1. Microplastic characterization

Fluorescently labelled polystyrene microspheres, suspended in water, were purchased from Polysciences Inc. (Warrington, PA) under the brand name, Fluoresbrite^®^ YG Microspheres, 3 µm in diameter (Cat. #: 18861-1) (1.68 × 10^9^ particles/mL), and 0.2 µm (Cat. #: 17151-10) (5.68 × 10^12^ particles/mL). According to the manufacturer, both have 441-nm excitation and 486-nm emission wavelengths. Stock solutions were protected from light and stored in the refrigerator (4 °C) until use. Working solutions of 10^7^ particles/mL (3 µm) and 10^9^ particles/mL (0.2 µm) were prepared using distilled water (DW).

### 2.2 Rotifer preparation

We employed euryhaline rotifers, *Brachionus plicatilis* (S-type), as larval feed. Rotifers were cultured with highly saturated, fatty acid-enriched *Chlorella* (Super *Chlorella* V12; *Chlorella* Industry) in 30-L batch cultures at 22 ppt (artificial seawater) and 25 °C with aeration. Daily concentrations of chlorella for rotifers were maintained as 2.5 × 10^6^ cells/mL. In order to maintain good water quality, 10-L aliquots of culture medium were replaced as needed. The required number of rotifers for each feeding regime was harvested, rinsed through a plankton net (45 µm mesh size), and transferred into three separate beakers (one containing only fresh seawater, and two beakers containing MPs or NPs, respectively) for an incubation period of 1 h. During this incubation period, rotifers ingested MPs or NPs, before being fed to bream larvae.

### 2.3. Larviculture

Fertilised eggs of red sea bream, *Pagrus major*, were purchased from ISC Co., Ltd. in Amakusa, Kumamoto Prefecture, Japan, and transported directly to the Fish and Ships Laboratory, Faculty of Fisheries, Nagasaki University. A total of 10,000 eggs were stocked in 100 L polycarbonate tanks with continuous aeration, at 35 ppt salinity. Water temperature was maintained at 19 °C using a room air conditioner. Fish eggs were acclimated and reared until hatching. Larviculture involved five treatments: control (no MPs or NPs), direct consumption of MPs in the water, direct consumption of NPs, indirect MP consumption via rotifer predation, and indirect NP consumption. Triplicate tanks were employed for each treatment (Fig. 1). When larvae opened their mouths (2 days post-hatch, dph) they were randomly transferred to experimental tanks at a density of 100 larvae per tank. Each 8-L rearing tank was filled with 35-ppt artificial seawater, and equipped with a centrally placed spherical aerator supplying air at 50 mL/min. Super *Chlorella* V12 was added daily to each experimental tank to achieve a density of 5 × 10^5^ cells/mL. For direct consumption experiments, the concentration of MPs or NPs in each rearing tank was maintained at 200 particles/mL (Seong et al., 2024), with a temperature of 19 ± 0.5 °C. The experiment was conducted for 12 days with a 12-h diurnal photoperiod (900 LJ 2100). Each tank was cleaned by siphon every 3 days to remove dead larvae, uneaten diet, and excrement.

**Figure 1:**
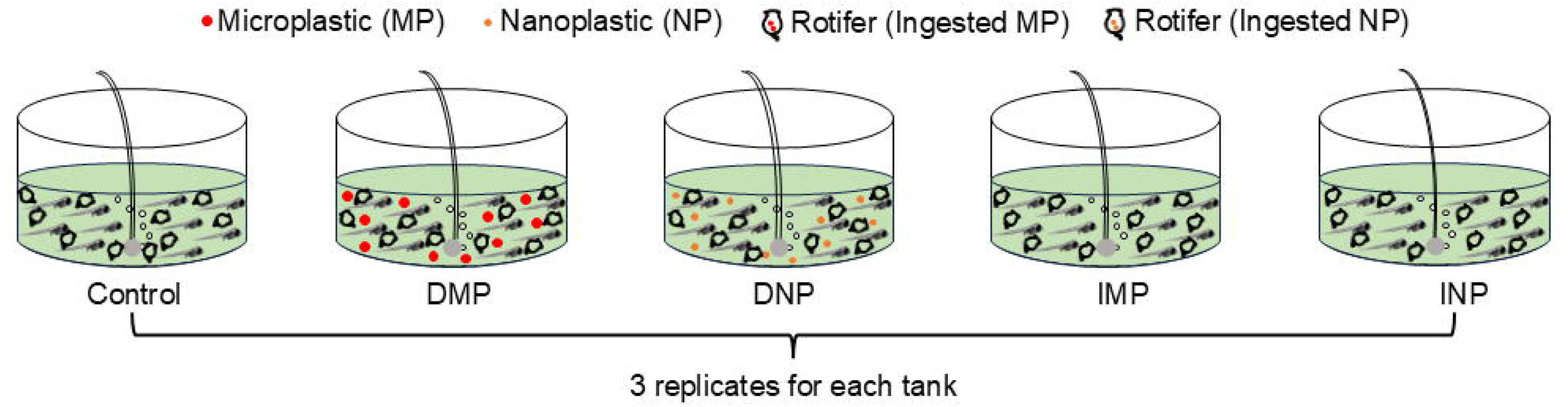
Schematic of experimental design for larviculture. Five experimental groups, control (no MPs or NPs), DMP (direct consumption of MPs in the water), DNP (direct consumption of NPs in the water), IMP (indirect MP consumption via rotifer predation), and INP (indirect NP consumption via rotifer predation). All experimental groups were fed with the same number of rotifers daily.

### 2.4. Feeding regimes

Two feeding regimes were employed to investigate direct and indirect consumption of MPs and NPs. Required numbers of rotifers were filtered and transferred into separate beakers. To ensure that the total MP or NP concentration via rotifer predation was similar to that for direct consumption (200 particles/mL), rotifer ingestion was monitored using fluorescence microscopy. Based on this quantification, rotifers were incubated at a concentration that enabled each individual to accumulate an average of 20 MPs or NPs, thereby having the same total particle count per mL as for direct exposure. After incubation, rotifers were filtered through a plankton net (45 µm mesh size), and fed to bream larvae at a density of 10 rotifers/mL. Rotifer density in all tanks was monitored daily and maintained across treatments by adding fresh rotifers when necessary.

### 2.5. Fish sampling

Sampling of larvae was conducted at 0, 2, 6, 9, 12 dph. At 0 and 2 dph, 30 larvae were collected randomly from the stock (100 L polycarbonate tank). For subsequent sampling (6, 9 and 12 dph), three larvae from each experiment tank were collected randomly. Larvae were anaesthetized using MS-222 (50 mg/L) and fixed in 4% neutral buffered formalin for measurement of total length (TL) and standard length (SL). Total and standard lengths were measured using a stereo microscope (Nikon, SMZ745T), and photographs were taken with an attached camera (WRAYCAM, NOA630). At the end of the experiment (12 dph), all remaining larvae were counted to determine the survival rate. For quantitative real-time PCR and enzyme assay, 30 larvae were collected from each experimental tank (divided equally for both analyses) and rinsed in distilled water. They were then placed in microtubes and stored at -80 °C until further processing. To evaluate the quality of fish eggs and hatched larvae, the hatching rate and survival activity index (SAI) of hatched larvae were also calculated. We placed 30 fertilized eggs in a 1-L beaker containing 500 mL of the same saline as for larviculture at 19 ± 0.5 °C in total darkness without aeration. Dead larvae were counted and removed every 24 h until total mortality, in order to estimate survival and resistance to starvation. Triplicate observations were used to calculate SAI according to the following equation:

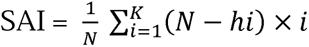

where N is the total number of examined larvae, hi is the cumulative mortality by the i-th day, and K is the number of days elapsed until all larvae died due to starvation.

### 2.6. Fluorescence image

Fish larvae were irradiated with blue light (excitation wavelength: 440 – 460 nm, fluorescence wavelength 500 nm) attached to a stereo microscope (Nikon, SMZ745T) to observe internalization and accumulation of MPs and NPs in larvae.

### 2.7. Antioxidant activity

Red sea bream larval SOD and CAT were measured in homogenates of 15 larvae. SOD activity was assayed using a commercial kit (Sigma-Aldrich, St. Louis, MO, USA; Cat. No. CS0009) by measuring inhibition of the reduction rate of cytochrome c by superoxide radical at 450 nm. One unit (U) of SOD activity is the amount of enzyme that resulted in 50% inhibition of cytochrome c. Meanwhile, CAT activity was assayed using a commercial kit (Abcam, Cambridge, UK; Cat. No. ab83464) by measuring residual H_2_O_2_ absorbance at 570 nm. Both results were expressed as U/mg protein.

### 2.8. Protein estimation

Protein content of red sea bream larvae was estimated using the Bradford assay. Bradford reagent (40 μL) and 160 μL of homogenate (15 larvae) were mixed, vortexed, and incubated at room temperature for 5 min. Bovine serum albumin (BSA) was used as standard, and absorbance was measured at 595 nm.

### 2.9. Gene expression

Total RNA was extracted from 15 fish from each replicate of each experimental tank. Larvae were homogenized in a NucleoSpin® RNA kit (Takara Bio Inc, Shiga, Japan) following the manufacturer’s protocol. RNA purity and concentration were checked spectrophotometrically at 230 nm, 260 nm, and 280 nm using a NanoDrop 2000 spectrophotometer (Thermo Fisher Scientific, Waltham, Massachusetts, USA). Subsequently, complementary DNA (cDNA) synthesis was performed using 1000 ng of RNA with a PrimeScript™ II 1st strand cDNA Synthesis Kit (Takara Bio Inc, Shiga, Japan), and was preserved at -20 °C until use. Quantitative real-time reverse transcription-polymerase chain reaction (qRT-PCR) was performed using 1 μL of cDNA template, 0.5 μL of forward and reverse primers (10 μM each), and 10 μL of TB Green Premix Ex Taq (2×) (Takara Bio Inc, Shiga, Japan) in a final volume of 20 μL. The assay was performed using a QuantStudio 1 Real-Time PCR System (Thermo Fisher Scientific, Waltham, Massachusetts, USA) with the following thermal cycling conditions: an initial denaturation at 95 °C for 30 s, followed by 40 cycles at 95 °C for 5 s and 60 °C for 30 s. Melting curve analysis was conducted to confirm specific product amplification, involving cycles at 95 °C for 15 s, 60 °C for 1 min, and 95 °C for 15 s following the protocol. Analyzed gene and primer sequences are listed in Table 1. The beta-actin (β-actin) gene was chosen as a reference due to its stable expression among experimental groups. Transcriptional levels were calculated using the 2^−ΔΔC^t method (Livak & Schmittgen, 2001). All experiments were performed in triplicate.

**Table 1:**
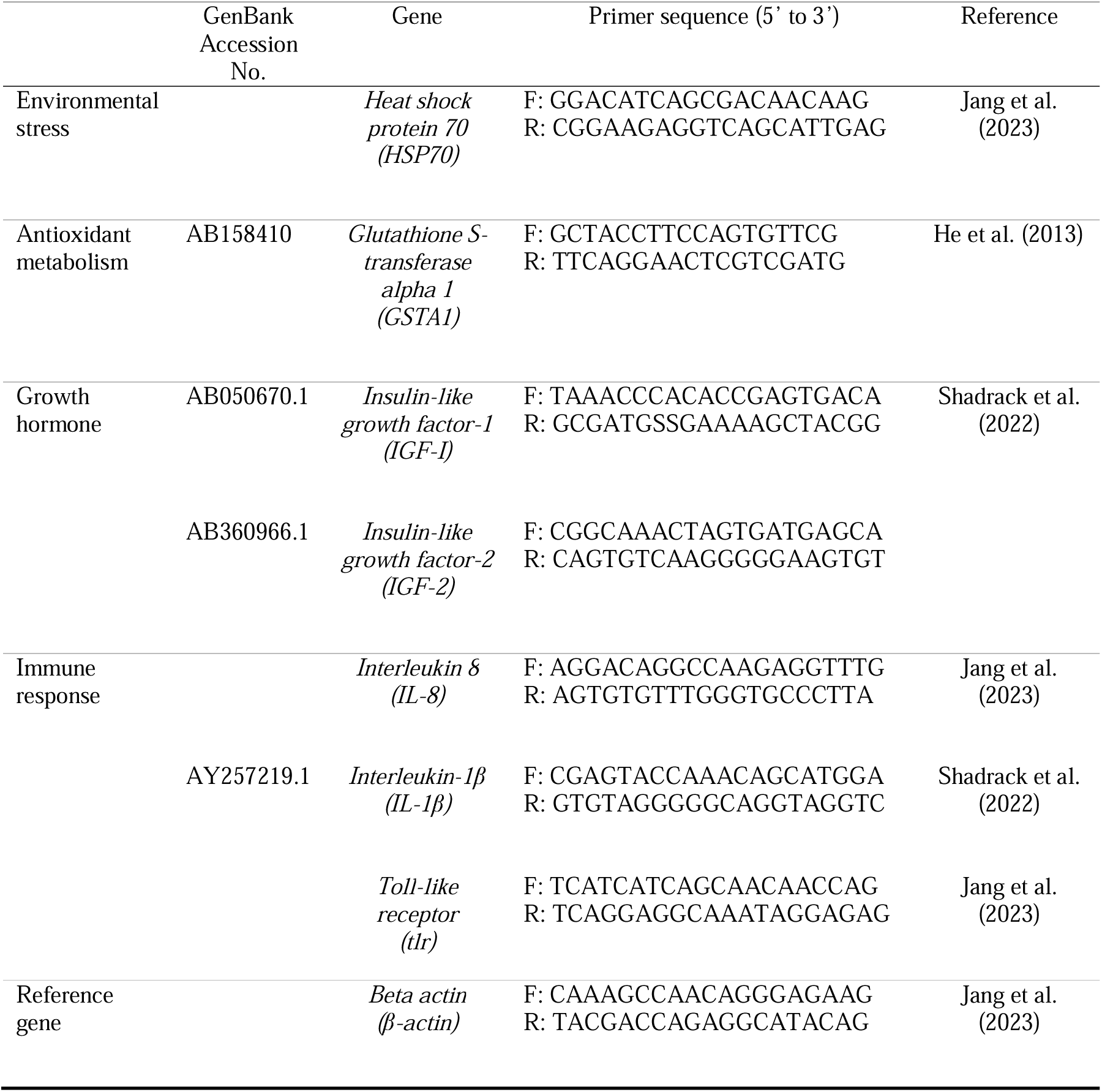
Primer sets employed in this study to assess biomarkers associated with environmental stress, antioxidant metabolism, growth hormone, and immune responses by larvae of *Pagrus major*.

### 2.10. Statistical analysis

The Shapiro-Wilk test was used to test for normality of survival, growth, and MP or NP effects on relative gene expression and antioxidant enzyme activity. Normally distributed data were compared by one-way analysis of variance (ANOVA). Prior to ANOVA, Levene’s test was used to confirm homogeneity of variances, and the Tukey HSD test was employed for post-hoc analysis. The Kruskal-Wallis test was performed for non-normally distributed data, and Dunn’s test was applied for post-hoc assessment if p<0.05. All statistical analyses were performed in R (v.4.3.3).

## 3. Results

### 3.1. Larviculture

Red sea bream eggs yielded a 96.7% hatching rate and hatched larvae from these eggs survived 12 days of starvation. The calculated larval survival activity index (SAI) was 51.2 ± 1.5. After 12 days of rearing, the survival rate of red sea bream larvae by both ingestion routes was lower than the control group. However, predation on rotifers fed NPs reduced the survival rate of red sea bream larvae significantly more than rotifers fed MPs (p < 0.01) (Fig. 2). However, total length and standard length of collected larvae did not differ significantly among treatment groups after exposure (Fig. 3).

**Figure 2:**
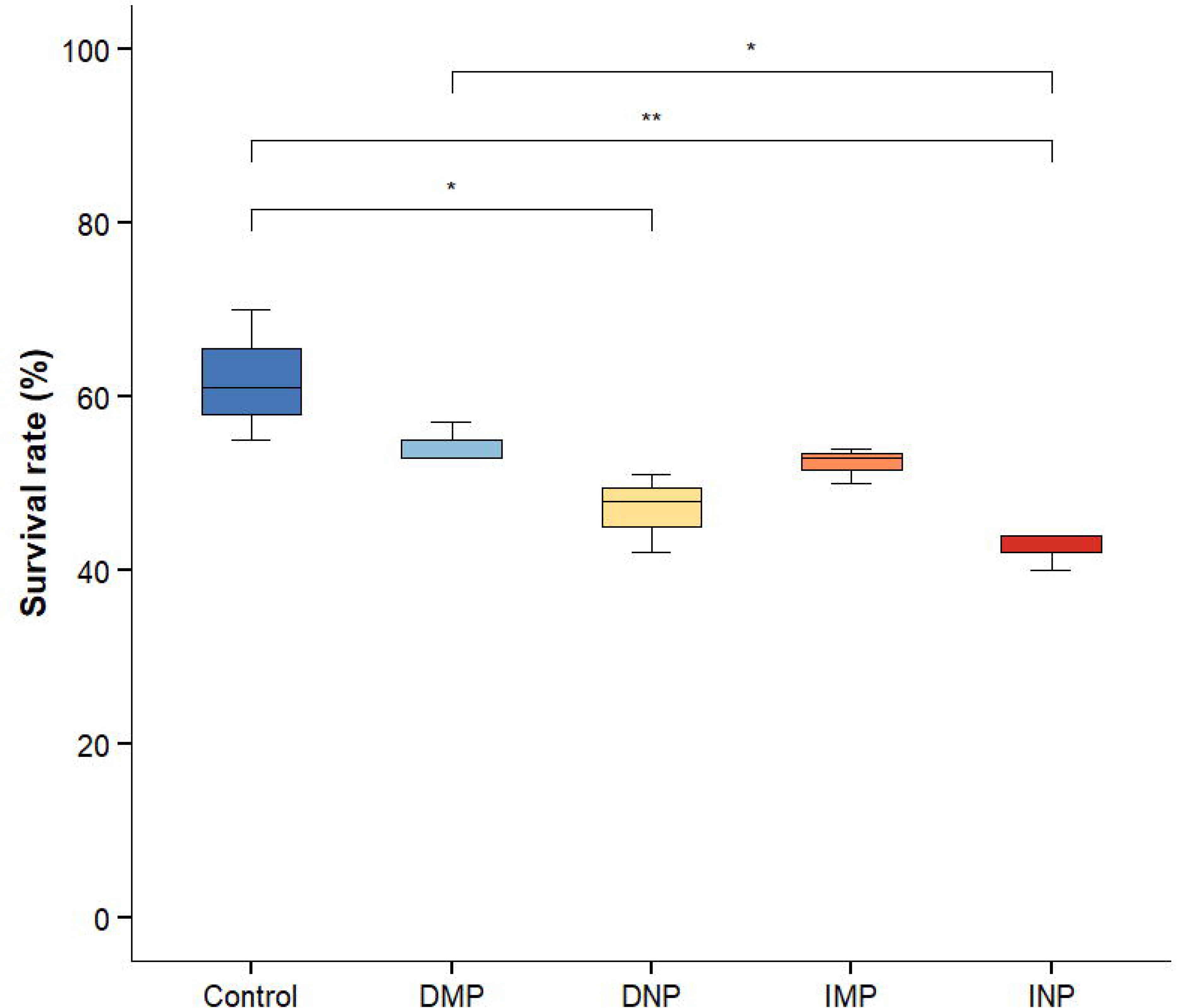
Box-and-whisker plots of the survival rate of red sea bream larvae for 12 days. Asterisks indicate significant differences (\**p*<0.05 and \*\**p*<0.01) by the Tukey post-hoc test.

**Figure 3:**
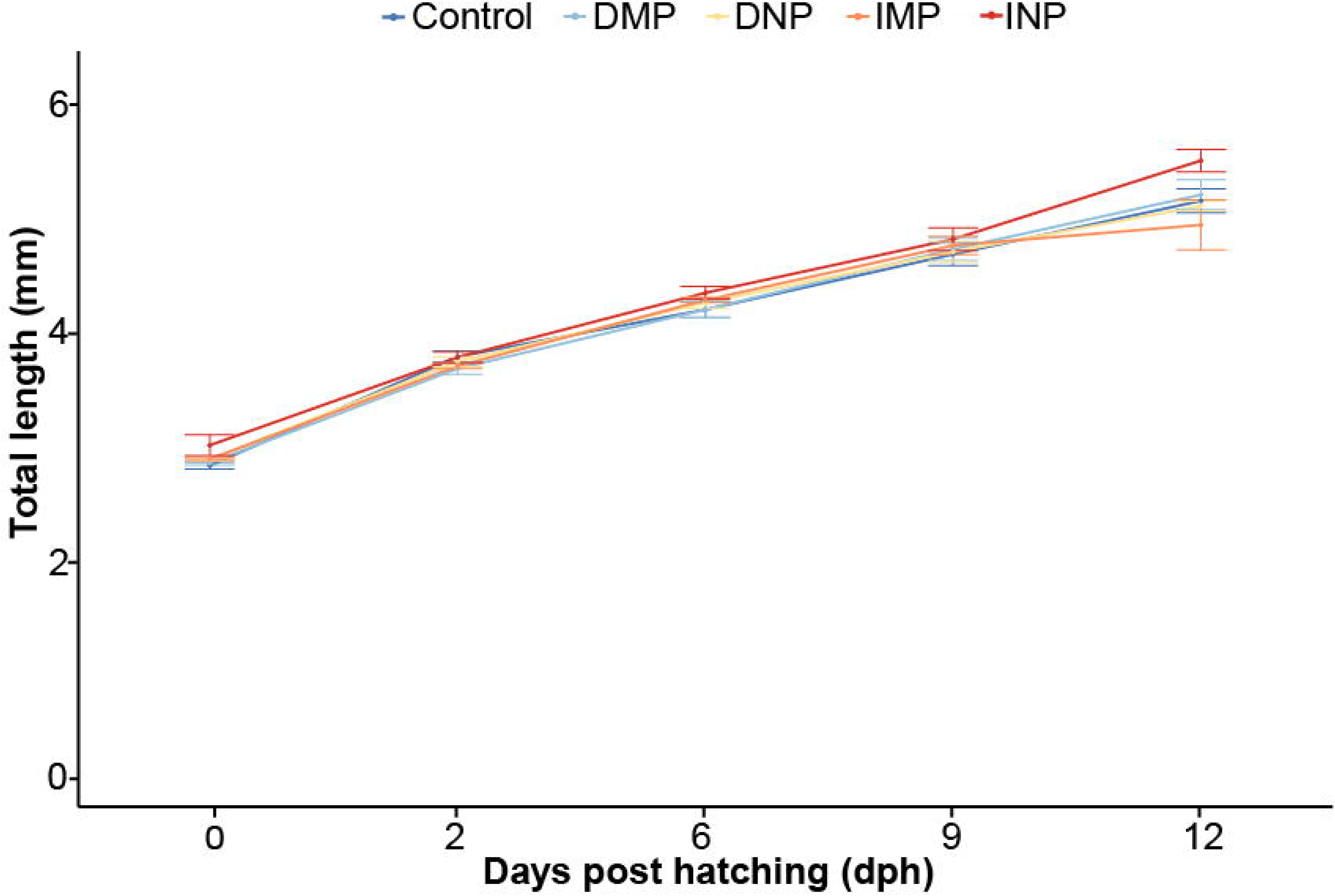
Total length of red sea bream larvae for 12 days. Points and error bars represent the means and standard deviations, respectively (n=3).

### 3.2. Antioxidant activities

To determine whether MPs or NPs induce oxidative stress, SOD and CAT activities were measured in bream larvae. After exposure to MPs or NPs, SOD activity levels of larvae were significantly increased (p<0.05) (Fig. 4). NPs via rotifer predation showed the highest SOD activity. Exposure to MPs or NPs increased bream CAT activity levels; however, differences among treatment groups were not significant (Fig. 4).

**Figure 4:**
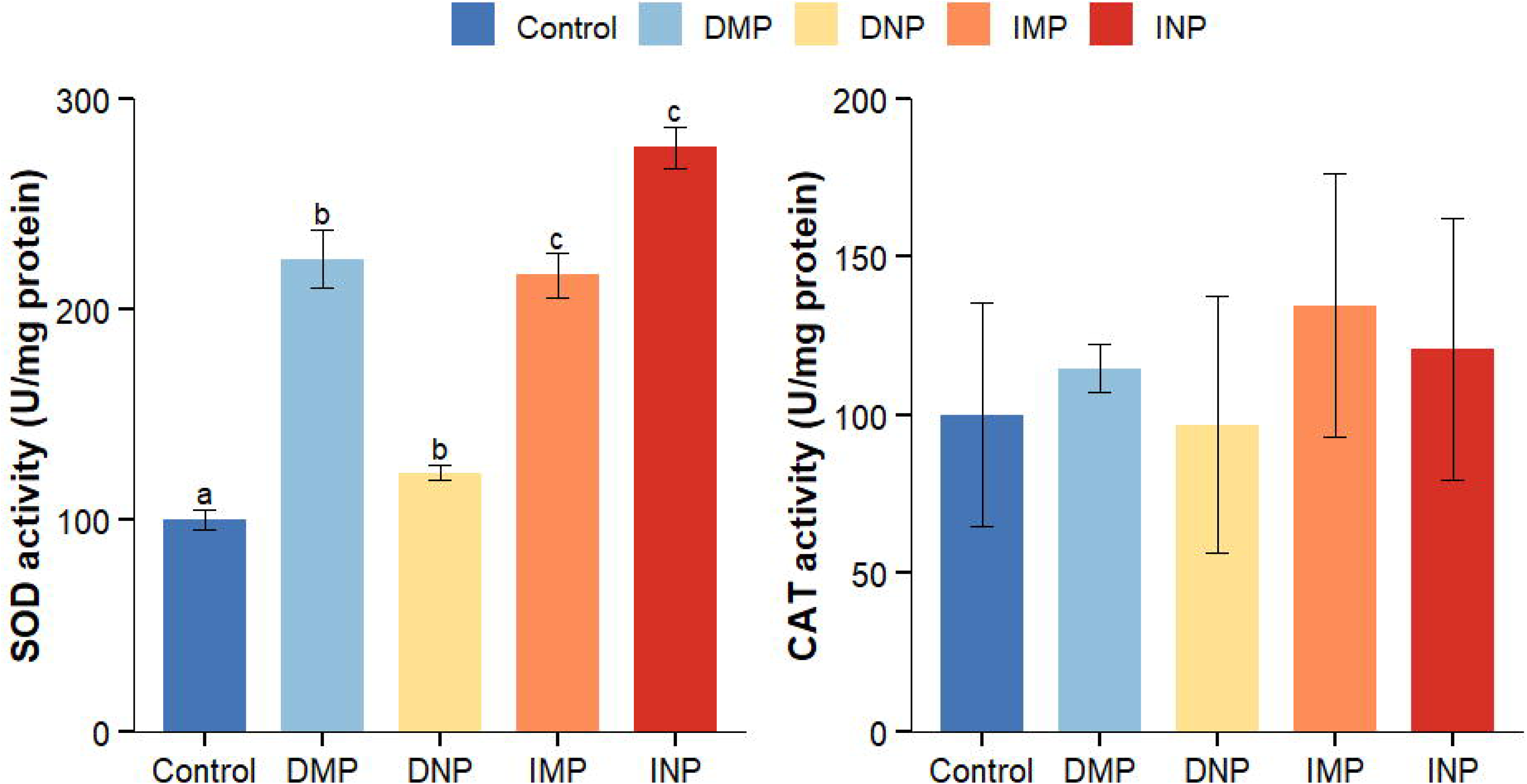
Antioxidant activity of red sea bream larvae induced under five plastic particle exposure treatments. Different superscript alphabet letters indicate significant differences among treatment groups (a > b > c, Tukey’s post-hoc test, *p*<0.05).

### 3.3. Relative gene expression

Expression of environmental stress, antioxidant, growth, and immune response genes of red sea bream larvae under MP and NP exposure is presented in Fig. 5. Overall, relative gene expression increased more significantly in response to rotifer predation than to direct consumption for all selected genes, except growth hormone. HSP70 gene expression was significantly downregulated in the MP group via rotifer predation compared to direct consumption (*p*<0.05, Fig. 5A). Expression of GSTA1 was also significantly upregulated in the MP group via rotifer predation compared to controls (*p*<0.05, Fig. 5B). For growth hormone expression, the highest expression of IGF-1 was recorded in the MP group via rotifer predation, but no significant differences were observed among experimental groups. However, expression of IGF-2 was significantly elevated in the MP group via rotifer predation compared to the control (*p*<0.05, Fig. 5C). Furthermore, expression of IL-1β, was significantly increased in the MP group via rotifer predation. However, there were no changes in IL-8 or tlr expression compared to controls (*p*<0.05, Fig. 5D).

**Figure 5:**
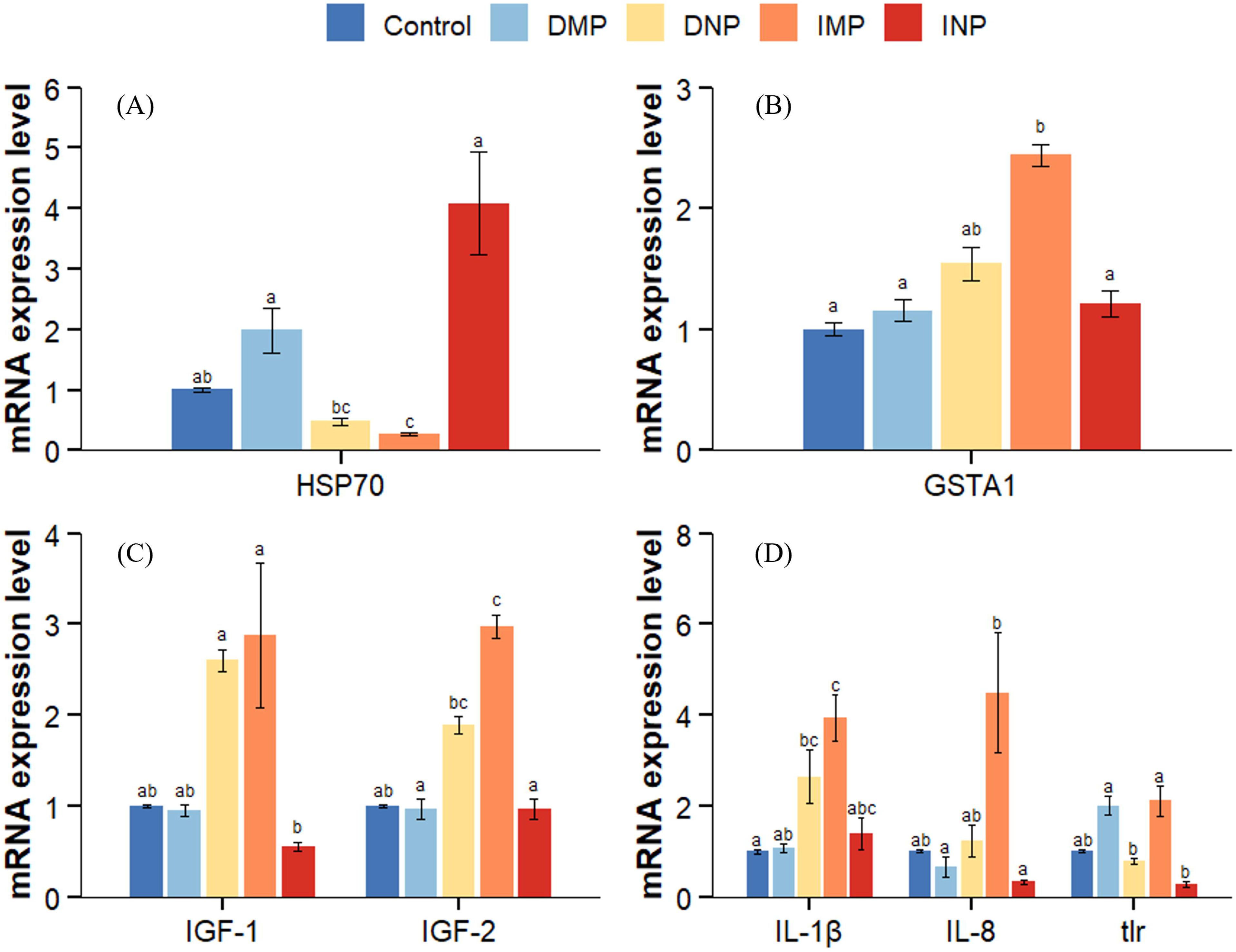
Plastic particle effects on relative gene expression in red sea bream larvae. Relative gene expression related to (A) environmental stress, (B) antioxidant response, (C) growth hormone, and (D) immune responses in red sea bream larvae. Different superscript alphabet letters indicate significant differences among treatment groups (a > b > c, Dunn’s post-hoc test, *p*<0.05).

## 4. Discussion

It has been reported that either micro- or nanoplastic ingestion can negatively impact aquatic animals, resulting in reduced growth and survival (Lu et al., 2016; Pelegrini et al., 2023). We observed significantly lower survival in both nanoplastic groups compared to corresponding microplastic groups. This is consistent with findings of Sendra et al. (2021), who reported reduced survival in zebrafish larvae exposed to polystyrene NPs (50 nm and 1 µm), even after only 24 h. However, Wen et al. (2018) found no significant effects in juvenile discus fish (*Symphysodon aequifasciatus)* after 30 days of polystyrene MP exposure, highlighting species-specific sensitivity and differences in exposure conditions. Although numerous studies have shown the capacity of some NPs to penetrate chorions, accumulate in tissues, and to impair fish locomotion and predation (Pitt et al., 2018; Zhang et al., 2021), NP biological impacts can vary depending on the test species and exposure pathways. In our study, no significant impairment of larval development was observed, consistent with previous findings in zebrafish larvae under similar exposure conditions (Lu et al., 2016; Feng et al., 2022). Additionally, Murray and Cowie (2011) demonstrated that plastic particles can accumulate and move through the food web. They detected polypropylene fibers in both prey and predators of Norway lobsters (*Nephrops norvegicus*), providing clear evidence of plastic movement across trophic levels. However, in our study, no fluorescent signals were visually confirmed in intestinal organs of larvae from rotifer predation due to extended storage (more than six months) before photography and dissection. Apparently, fluorescence degrades over time in rotifer digestive tracts, as in tissue (Zhang et al., 2025), and further confirmation through histological analysis is necessary to verify internal localization and retention of plastic particles in internal organs.

Micro- and nanoplastics are increasingly recognized as environmental stressors in aquatic organisms (Santos et al., 2021; Abdel-Latif et al., 2022). One key mechanism by which these particles exert toxicity is by stimulating overproduction of ROS, overwhelming the antioxidant defense system. The SOD-CAT system usually serves as the first line of defense against oxidative stress (Romano et al., 2020). SOD catalyzes conversion of superoxide radicals into H_2_O_2_ by SOD, which is subsequently degraded by CAT into water and oxygen. In our study, SOD activity was significantly elevated in all exposure groups, indicating the onset of oxidative stress. Similar findings were reported in zebrafish and Nile tilapia (*Oreochromis niloticus*) exposed respectively to polystyrene MPs of different sizes (5 µm and 0.1 µm) (Lu et al., 2016; Ding et al., 2018). CAT activity was also elevated, but not significantly in any group, which may be due to the relatively short exposure duration. In contrast to these findings, Kang et al. (2021) observed a decrease in SOD and CAT activities in guts of marine medaka (*Oryzias melastigma*) after 14 days of polystyrene MP exposure. Additionally, Zhang et al. (2021) revealed increased SOD activity accompanied by elevated malondialdehyde (MDA) levels in intestines of silver carp (*Hypophthalmichthys molitrix*), indicating that while antioxidant defenses were activated, they were insufficient to prevent oxidative damage. These findings collectively suggest that while MPs or NPs can activate early antioxidant responses, effectiveness of the defense system may vary depending on exposure duration, species, or plastic characteristics.

In addition to the above antioxidant enzymes, GSTA1 is vital for detoxification processes in various organisms. In this study, GSTA1 expression was significantly increased only after MP-rotifer predation, suggesting that trophic transfer induces a more pronounced oxidative stress response than direct consumption. This increase likely reflects activation of the glutathione-dependent antioxidant defense system in response to MPs. Similarly, Wen et al. (2018) observed elevated glutathione (GSH) levels in *S. aequifasciatus* exposed to 500 µg/L MPs, indicating induction of redox-regulating mechanisms. In contrast, Lu et al. (2018) found decreased GSH content in zebrafish exposed to a mixture of MPs and cadmium (Cd), suggesting that co-exposure may overwhelm antioxidant defenses. Overall, the exclusive and significant elevation of GSTA1 in the MP group via rotifer predation highlights enhancement of oxidative stress specifically associated with trophic transfer of MPs in red sea bream larvae.

Heat shock proteins (HSPs) are highly conserved molecular chaperones involved in protein folding and remodeling, particularly under stress, to maintain cellular homeostasis and promote cell repair (Zhang et al., 2014). Here, we observed a significant decrease in HSP70 expression in the MP group via rotifer predation, suggesting that oxidative stress by this exposure route may not be sufficient to trigger a detectable HSP70 response. A similar result occurred in zebrafish embryos exposed to polystyrene NPs, indicating suppression or bypassing of the classical stress-response pathway (Martin-Folgar et al., 2023). Contrary to our finding, Das et al. (2023) reported upregulation of HSP70 in Nile tilapia exposed to polystyrene MPs, reflecting a typical cellular stress and xenobiotic detoxification response. Increased HSP70 mRNA expression can enhance immune and protective responses under environmental stress (Fu et al., 2011; Zhang et al., 2014). Therefore, downregulation observed in our study may reflect size- or exposure pathway-dependent suppression of this stress response, or delayed transcriptional activation in red sea bream larvae.

Furthermore, significant growth hormone expression was only recorded in mRNA expression of IGF-2, which was elevated after MP-rotifer predation. Notably, we observed no reduction in somatic growth of red sea bream larvae after 12 days of exposure to polystyrene MPs or NPs via either consumption route, which suggests that trophic transfer exposure may not impair early development in this species. However, Barboza et al. (2018) and Pitt et al. (2018) documented suppression of growth hormone signaling leading to growth retardation in gilthead sea bream (*Sparus aurata*) and zebrafish under NP exposure, which conflicts with our findings. Similarly, Romano et al. (2020) found that polyvinyl chloride MP exposure significantly reduced growth in goldfish. Upregulation of IGF genes observed in our study may reflect compensatory growth-promoting mechanisms in response to MPs or may indicate species-specific or exposure route differences in physiological impacts of MPs.

Fish immune systems can be impacted by MPs or NPs through damage to the gut due to cytokine production, as well as by inflammation of the gut (Bhagat et al., 2020). Once ingested, plastic particles may interact with intestinal tissues to enter the bloodstream, potentially modulating immune function (Hirt & Body-Malapel, 2020). In this study, a significant increase in IL-1β gene expression was observed only in the MP group via rotifer predation, indicating immune system activation and potential inflammatory responses. Similarly, Choi et al. (2018) observed elevated inflammatory genes in tilapia, and Jin et al. (2018) found a similar response in zebrafish after polystyrene MP exposure. However, no significant changes were observed for IL-8 and tlr expression in any group, suggesting that the immune response may be limited to specific pathways or cytokines, potentially depending on plastic particle size or exposure routes. In contrast, Mazurais et al. (2015) reported no significant changes in IL-1β expression in European sea bass larvae after 43 days of exposure to polyethylene microbeads, highlighting variability in immune responses between species and plastic types. Therefore, selective upregulation of IL-1β in larvae fed MP-rotifers, suggests that trophic exposure to MPs may be sufficient to trigger an early pro-inflammatory response, while other immune pathways remain unaffected at this stage of development.

## 5. Conclusions

In conclusion, this study provides compelling evidence that MPs and NPs, even in early trophic levels, can significantly compromise survival and physiological homeostasis of marine fish larvae. While somatic growth was not affected during 12 days of exposure, notable molecular and biochemical alterations were observed, indicating sublethal stress responses that may portend long-term developmental impairment. Specifically, survival was significantly lower in both nanoplastic groups than in corresponding microplastic groups, suggesting higher toxicity associated with smaller particle sizes. Nonetheless, survival of bream larvae that consumed NPs via rotifer predation was significantly lower than those that consumed NPs directly. Despite a lack of observable developmental abnormalities, elevated expression of growth-related genes (IGF-1 and IGF-2), especially in the MP group via rotifer predation, may point to compensatory growth signaling mechanisms rather than to true physiological stability. Elevated SOD activity among all exposure groups indicates oxidative stress induction, whereas exclusive upregulation of GSTA1 in the MP group via rotifer predation suggests enhanced activation of glutathione-mediated detoxification. Interestingly, HSP70 was downregulated under the same conditions, pointing to potential suppression or a delay in stress response activation. The selective increase in IL-1β expression in the MP group via rotifer predation further indicates early-stage immune activation specific to trophic exposure.

Overall, these findings suggest that trophic transfer induces more pronounced biological effects than direct plastic consumption. This underscores the need to consider ingestion route, particle size, and life stage when assessing MP or NP toxicity. Future studies should investigate long-term and transgenerational effects of both MPs and NPs under environmentally relevant concentrations to better evaluate ecological risks and potential food-web-level impacts in marine ecosystems.

## CRediT authorship contribution statement

**Siti Syazwani Azmi:** Conceptualization, Formal analysis, Investigation, Writing – original draft; **Ozan Oktay:** Conceptualization, Formal analysis, Investigation; **Hee-Jin Kim:** Conceptualization, Investigation, Writing – review & editing; **Hisayuki Nakatani:** Investigation, Writing – review & editing; **Mitsuharu Yagi:** Supervision, Project administration, Writing – review & editing.

## Data availability

Data will be made available on request.

